# Learning the Language of the Microbiome with Transformers

**DOI:** 10.64898/2026.05.02.722381

**Authors:** Neythen J Treloar, Saif Ur-Rehman, Jenny Yang

**Affiliations:** Outpost Bio

## Abstract

Self-supervised pretraining has become central to biological machine learning, yet microbiome data remains comparatively underexplored in terms of both modeling approaches and evaluation frameworks. To address this gap, we present Atlas, a pretraining dataset of 539,308 microbiome datapoints from the MGnify database. Using Atlas, we train the Waypoint family of microbiome foundation models: a series of GPT-2 style causal language models ranging from 6M to 170M parameters. We also introduce Compass, a curated benchmark of eight predictive tasks spanning biome classification, drug-microbiome interactions, drug degradation, and infant gut development. Using this benchmark, we compare the performance of Waypoint models against classical baselines and the existing Microbial General Model (MGM) foundation model. Our results show that pretraining leads to consistent and significant improvements in downstream task performance, that both dataset scale and tokenization strategy impacting model quality, that pretraining is essential for achieving favorable scaling behavior and that representations learned during pretraining generalise between microbiome domains. Furthermore, pretrained transformer models begin to reliably outperform classical methods once training data exceeds roughly 10,000 examples - a threshold that is attainable for modern microbiome studies. Finally, we demonstrate that the Waypoint models achieve state-of-the-art performance among microbiome foundation models. Overall, our work highlights the importance of large-scale self-supervised pretraining and tokenisation choices in this domain and establishes Atlas, Compass, and the Waypoint models as valuable resources for the research community in this emerging field.

## 1 Introduction

Microbial communities, from the human gut to marine sediments and engineered systems, have emerged as a powerful variable in human health, disease, and environmental biology. Advances in high-throughput sequencing have made it possible to profile microbial communities at scale, generating large repositories of compositional abundance data across diverse ecological contexts. Yet translating this wealth of data into actionable biological insight remains a significant challenge, owing to the high dimensionality, sparsity, and compositional nature of microbiome profiles, as well as the heterogeneity in sequencing and taxonomic assignment pipelines.

Much recent progress in machine learning has been driven by the use of self-supervised pretraining to learn meaningful representations, especially in language modelling. These techniques have also been applied to biological data, giving rise to DNA language models [1, 2, 3, 4, 5], protein language models [6, 7], and whole-genome models [8, 9]. This naturally raises the question of whether the architectures and training paradigms that succeed in natural language and molecular biology transfer effectively to microbiomes and this is an emerging frontier [10, 11, 12]. As this progresses it is also becoming increasingly important to evaluate which language modelling techniques are truly applicable, and to establish benchmarks that can guide the field as it develops.

The foundation model paradigm is particularly attractive for microbiome research, where labelled datasets are typically small and task-specific, but large collections of unlabelled compositional profiles are publicly available. Several recent efforts have begun to treat taxonomic abundance profiles as sequences amenable to language model pretraining [10, 11, 12]. However, these approaches differ in their tokenisation strategies, pretraining corpora, and architectural choices, and systematic evaluation of how these design decisions affect downstream performance has been limited. In particular, the effect of model scale on microbiome representation learning, a question extensively studied in language [13, 14] and protein [15] modelling, remains poorly understood. Equally, the field lacks standardised benchmarks for comparing models across the range of biologically meaningful prediction tasks that microbiome data can support.

In this paper, we take first steps toward addressing these gaps. We assemble and release a large-scale pretraining corpus, named Atlas, of 539,308 microbiome datapoints from the MGnify database [16], spanning amplicon, shotgun metagenomic, and metagenomic assembly modalities across a broad range of environments. We introduce a fallback tokenisation strategy that preserves taxonomic information for samples where genus level assignments are unavailable, and show that this strategy, combined with a larger pre-training corpus, improves on the prior state-of-the-art [10]. We also present Compass, a curated benchmark of eight predictive tasks spanning biome classification, drug-microbiome interactions, drug degradation, and infant gut development, designed to provide a comprehensive and reproducible evaluation framework for microbiome foundation models. Finally, we pretrain the Waypoint models, a series of GPT-2 [17] style causal language models ranging from 6M to 170M parameters, and evaluate each on Compass, along-side classical baselines and the existing MGM foundation model [10]. Our results demonstrate that pretraining consistently and significantly improves downstream performance, enables larger models to be used effectively and that representations learned druing pretraining generalise across microbiome domains. We also demonstrate that for downstream finetuning the amount of task-specific data is a key determinant of model quality. Finally, Waypoint models achieve state-of-the-art performance among microbiome foundation models, with dataset scale and tokenisation strategy emerging as important drivers of model quality.

## 2 Results

### 2.1 Atlas: the microbiome pretraining dataset

We assembled a large-scale pretraining corpus by systematically scraping taxonomic abundance data from the MGnify database [16] across all available pipeline versions (v1.0–v5.0) and four sequencing modalities: amplicon 16S rRNA, whole-genome shotgun metagenomic, metagenomic assembly, and metatranscriptomic (see Methods for details). The resulting raw collection spans over 4,100 unique studies, representing one of the largest aggregations of publicly available microbiome data to date.

Before tokenisation we quality filter the dataset, following a procedure used previously in related work [10], by removing any taxa with relative abundance lower than 0.0001 and subsequently removing any samples with less than 10 taxa. Following quality filtering, the dataset comprises 539,308 microbiome datapoints drawn from a broad range of environments including marine, freshwater, terrestrial, host-associated, and engineered ecosystems, reflecting the ecological diversity of the underlying MGnify resource.

Previous approaches have tokenised microbiomes at the genus level [10, 11], but other work has argued for the advantages of tokenising at the species level [12]. Tokenising at the species level can improve taxonomic precision; however, for many Amplicon sequence variants (ASVs) lower-level taxonomic assignments cannot be made. This is particularly relevant for amplicon sequencing, where the 16S rRNA marker often lacks the resolution needed to assign an ASV at the species level. To address this limitation, we introduce a fallback tokenisation approach in which a target level of taxonomic specificity is defined. Classifications more specific than the target rank are collapsed to that rank, while taxa that cannot be assigned at the target rank are represented using the most specific higher-rank classification available. To determine whether to tokenise at the genus or species level, we analysed the dataset using both tokenisation strategies. We found minimal difference in the distribution of sequence lengths and the number of total tokens between the two approaches (Table 1, Figure S1) and so we decided to move forward with genus level tokenisation with fallback (see Methods for details).

**Table 1:**
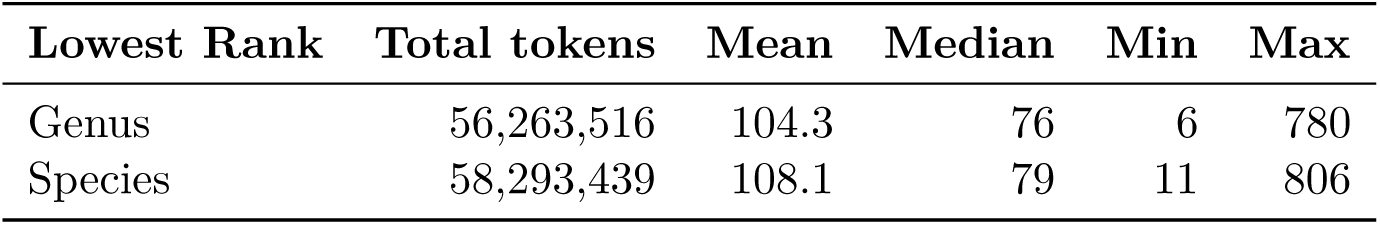
Tokenised corpus statistics at genus and species resolution. Sequence length is measured as tokens per sample. Mean, Median, Min and Max refer to sequence length across the dataset.

### 2.2 The Waypoint series of models: scaling microbiome foundation models

To investigate the effect of model scale on microbiome representation learning, we pretrained a series of GPT-2–style causal language models ranging from 6M to 170M parameters (not including token and positional embeddings) See Table 2 for architectural parameters and Methods for further details. Models were scaled by jointly increasing the hidden dimension and the number of transformer layers, while holding the per-head dimension fixed at 64 throughout (i.e. the number of attention heads grows in proportion to the hidden dimension). The resulting series spans roughly one order of magnitude in parameter count (6M - 170M). We also included an alternate 6M model matching the MGM [10] architecture (6M-MGM) enabling us to understand the performance gain from using a larger pretraining dataset and fallback tokenisation, and a 85M-gpt2-small architecture corresponding to the GPT-2 small model [17]. All models share the same tokeniser, maximum context length (512), and pretraining procedure, so that differences in downstream performance can be attributed to model capacity alone.

**Table 2:** Model configurations used in the scaling study. Layers denote the number of transformer blocks, hidden dimension the size of the model’s internal representations, and heads the number of attention heads per layer.

| Model | Layers | Hidden dimension | Number of Heads |
| --- | --- | --- | --- |
| 6M-MGM | 8 | 256 | 8 |
| 6M | 8 | 256 | 4 |
| 10M | 8 | 320 | 5 |
| 18M | 10 | 384 | 6 |
| 29M | 12 | 448 | 7 |
| 45M | 14 | 512 | 8 |
| 79M | 16 | 640 | 10 |
| 85M-gpt2-small | 12 | 768 | 12 |
| 170M | 24 | 768 | 12 |

### 2.3 Larger scale waypoint models show improved pretraining performance

We pretrained all models with an autoregressive next token prediction objective (see Methods for details). Pretraining evaluation loss curves for all nine models are shown in Figure 1A. All models converge from a shared initial loss of 5.0, with larger models achieving consistently lower evaluation loss throughout training. The two 6M-parameter models (6M and 6M-MGM) produce almost indistinguishable loss curves, demonstrating that the difference in attention head count between the two architectures has a minimal effect on pretraining performance. Scaling up model capacity yields consistent improvements, with eval loss decreasing monotonically with model size (Figure 1B). Interestingly the 79m and 85m-gpt-small models show very similar loss curves, with the difference in final loss being primarily due to the 79m model hitting the early stopping condition early and exiting training, despite being slightly different in size and architecture. Larger models also converge in fewer steps, while smaller models continue to improve slowly, suggesting that the smaller models have not fully converged and further gains remain available with extended training.

**Figure 1:**
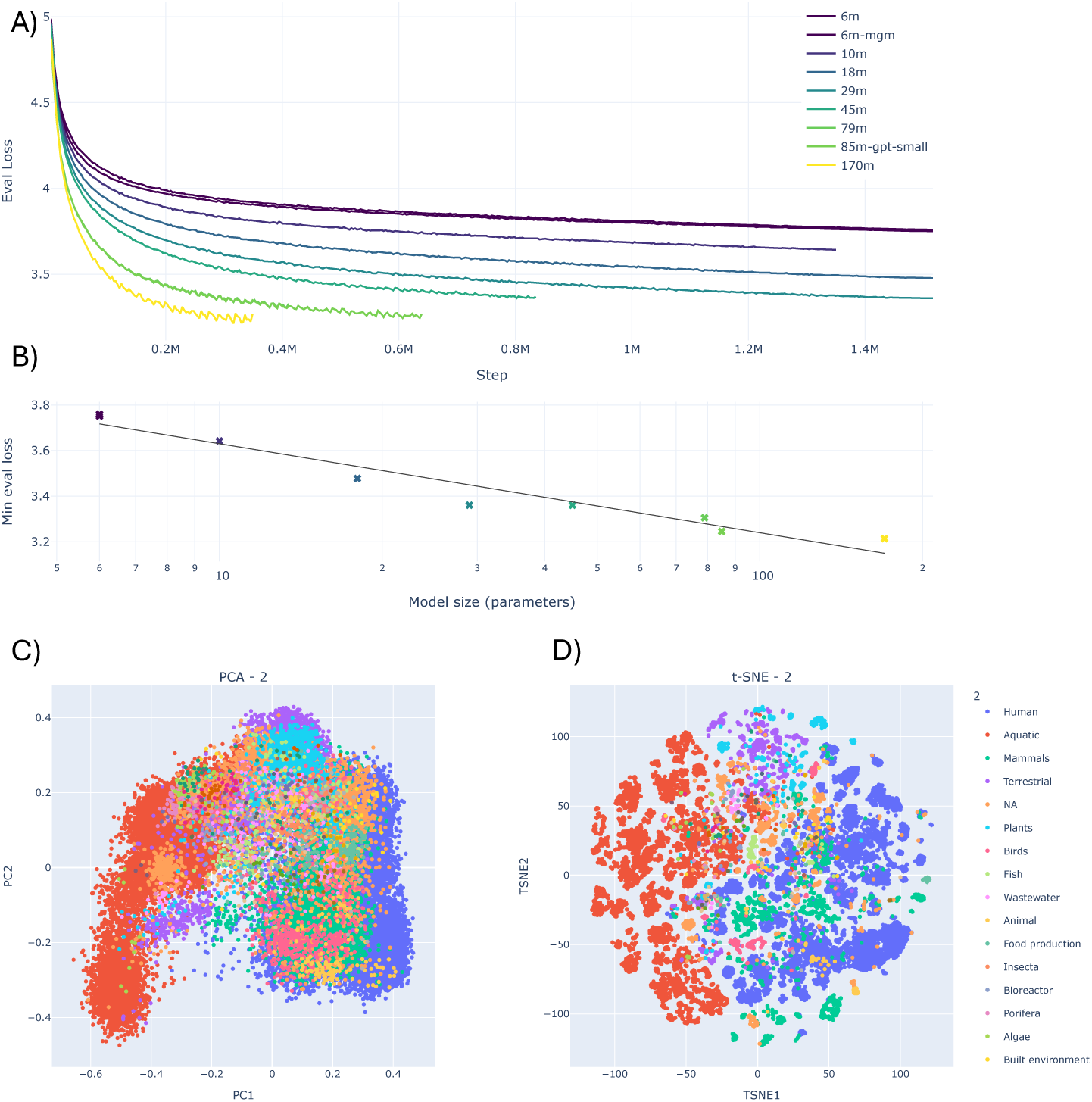
A) Pretraining evaluation loss curves for models with 6 - 170 million parameters. Larger models converge to lower evaluation loss. Some models exited training early because early stopping was used with a patience of 10. B) The minimum evaluation loss decreases linearly with logarithmic model size. C-D) PCA and t-SNE plots of mean pooled model embeddings coloured according to the second level of biome hierarchy.

Microbiome embeddings were generated from the 170M model by mean pooling over output token embeddings. In both PCA (Figure 1C) and t-SNE (Figure 1D) projections of these embeddings we see clear structure with respect to the second level of the biome hierarchy (for all other levels see Figure S2).

### 2.4 Compass: the microbiome benchmark

To systematically evaluate these pretrained models we introduce Compass, a curated benchmark of eight predictive tasks spanning diverse gut and environmental microbiome tasks (see Methods for details). The benchmark is designed to evaluate the ability of models to extract biologically meaningful information from compositional microbiome data. For now we focus primarily on gut microbiome problems, but this can be expanded upon in the future.

**Table 3:** Compass task summary. Classification tasks (task 1-5 and 7-8) are scored by macro-averaged F_1_; the regression task (task 6) by *R*^2^.

| Task | Name | Dataset | Type | Metric | N Training Points |
| --- | --- | --- | --- | --- | --- |
| 1 | Biome classification | [16] | Classification | $F_1$-macro | 33,121 |
| 2 | Gut biome classification | [16] | Classification | $F_1$-macro | 10,016 |
| 3 | SDC classification | [18] | Classification | $F_1$-macro | 2,725 |
| 4 | Drug vs. non-drug | [18] | Classification | $F_1$-macro | 3,168 |
| 5 | Drug class (ATC) | [18] | Classification | $F_1$-macro | 1,425 |
| 6 | Drug degradation rate | [19] | Regression | $R^2$ | 9,282 |
| 7 | Infant age classification | [20] | Classification | $F_1$-macro | 1,625 |
| 8 | Delivery mode classification | [20] | Classification | $F_1$-macro | 1,625 |

The eight tasks are drawn from four independent datasets (Table 3). The MGnify dataset [16] contributes two biome classification tasks: task 1 requires predicting the environmental origin of a microbiome sample at up to five levels of a biological ontology hierarchy (e.g., Environmental Aquatic Marine), while task 2 restricts this to classification of data points from gut microbiomes. The Shi et al. dataset, derived from a 16S rRNA amplicon study of drug–microbiome interactions in stool derived communities [18], contributes three tasks: task 3 requires classifying the sample from which a drug perturbed community originated from; task 4 is a binary classification of whether a sample received a drug or was a control; and task 5 requires predicting the ATC drug class that was applied to a community from the resulting composition. The Mastrorilli et al. dataset [19] contributes task 6, a regression task predicting drug degradation rate from community composition and drug identity. Finally, the Roswall et al. infant microbiome dataset [20] contributes task 7, predicting the age of an infant from a gut microbiome sample, and task 8, predicting delivery mode (vaginal vs. cesarean section).

Tasks 1–5 and 7–8 are classification tasks scored by macro-averaged F1 across classes, averaged across target columns where multiple targets are present (tasks 1 and 2). This scoring scheme weights classes equally regardless of prevalence, reflecting the benchmark’s aim to assess generalisation across the full label space rather than rewarding majority-class prediction. Task 6 is a regression task scored by the coefficient of determination (*R*^2^), clipped to be greater or equal to zero to prevent large negative values from skewing the overall score. To prevent datasets represented by multiple benchmark tasks from disproportionately influencing aggregate performance, we used dataset-balanced averaging. For each model and replicate, task scores were first averaged within each source dataset and the mean of these four averages was taken as the final dataset score (see Methods for details).

### 2.5 Pretrained transformers outperform non pretrained transformers and baseline models

We evaluated our pretrained Waypoint models against classical baselines and the current state of the art in microbiome foundation models. First, as classical baselines, we included simple multi layer perceptron (MLP), linear models, random forest models and XGBoost operating on bag-of-taxa abundance matrices (see Methods for details). All Waypoint and baseline models were trained with class weighted losses to handle class imbalance and simple hyperparameter optimisation was done for each baseline (see Methods). The sequencing data in each dataset used in the benchmark was processed differently by different authors, meaning that coverage of downstream taxa by the pretraining vocabulary varied substantially across datasets (Table 4). This variation in vocabulary overlap directly motivated evaluating the classical baselines both with and without filtering of taxa absent from the pretraining vocabulary (the filtered versions denoted “no UNK”), where the latter removes all out-of-vocabulary taxa to enable a fairer comparison to the Waypoint models operating under a fixed token set. Note that even in the no UNK setting the baseline models still have an information advantage as they are given the precise relative abundances of each taxon, whereas the Waypoint models just see rank ordered lists of taxa. Second, we compared against MGM, an existing microbiome foundation model pretrained on 260K samples from MGnify using a causal language modelling objective [10]. Our 6M-MGM model uses the same GPT-2 architecture as MGM and thus provides a direct assesment of the effect of our larger pretraining corpus and fallback tokenisation scheme with the prior state of the art. We also attempted to include comparisons with two additional microbiome foundation models [11, 12], but to our knowledge these models have not been made publicly available. For each transformer architecture, we additionally evaluated non-pretrained variants by reinitialising weights prior to benchmarking.

**Table 4:** Coverage of unique (the proportion of tokens in the vocabulary that are in the pretraining data) and total tokens (the proportion of total tokens across the dataset that are in the pretraining data) for benchmark datasets.

| Dataset | Tasks | Unique Token Coverage | Total Token Coverage |
| --- | --- | --- | --- |
| MGnify-biomes | 1–2 | 100% (complete) | 100% (complete) |
| Shi et al. | 3–5 | 73.2% (139/190) | 77.4% (53,884/69,659) |
| Mastorilli et al. | 6 | 95.9% (231/241) | 93.5% (1,576,034/1,686,268) |
| Roswall et al. | 7–8 | 60.5% (353/583) | 59.8% (52,018/87,027) |

Figure 2 shows the overall Compass score for each model and baseline, with replicates shown as individual points. In terms of the overall benchmark score the MLP baseline is not competitive, the Linear and Random Forest models are competitive with our non-pretrained transformers and the XG-Boost models are competitive with our pretrained transformers. We see that all of our pretrained transformers outperform XGBoost apart from 6M-MGM. Further, we see that for every baseline model the no UNK variant underperforms the variant that can see all the tokens, this highlights that having full knowledge of the taxa is important for good performance and implies that the token mismatch between pretraining and benchmarking is likely having a deleterious effect on the performance of the Waypoint models. We also see that pretraining consistently and substantially improves performance over both non-pretrained counterparts, and all pretrained Waypoint models outperform the MGM model, which itself outperforms all non-pretrained transformers. Additionally, 6M-MGM outperforms the original MGM despite sharing the same architecture, indicating that the larger pretraining dataset and the more informative input sequences enabled by fallback tokenisation contribute to improved performance. The effect of the improved tokenisation scheme is further supported by comparing the non-pretrained MGM and 6M-MGM models, where the only difference is the tokenisation scheme; the superior performance of 6M-MGM in this setting highlights the importance of tokenisation as a key modelling decision. Non-pretrained transformers exhibit substantially higher variance across replicates, indicating that these architectures are difficult to train from random initialisation on the relatively small per-task datasets.

**Figure 2:**
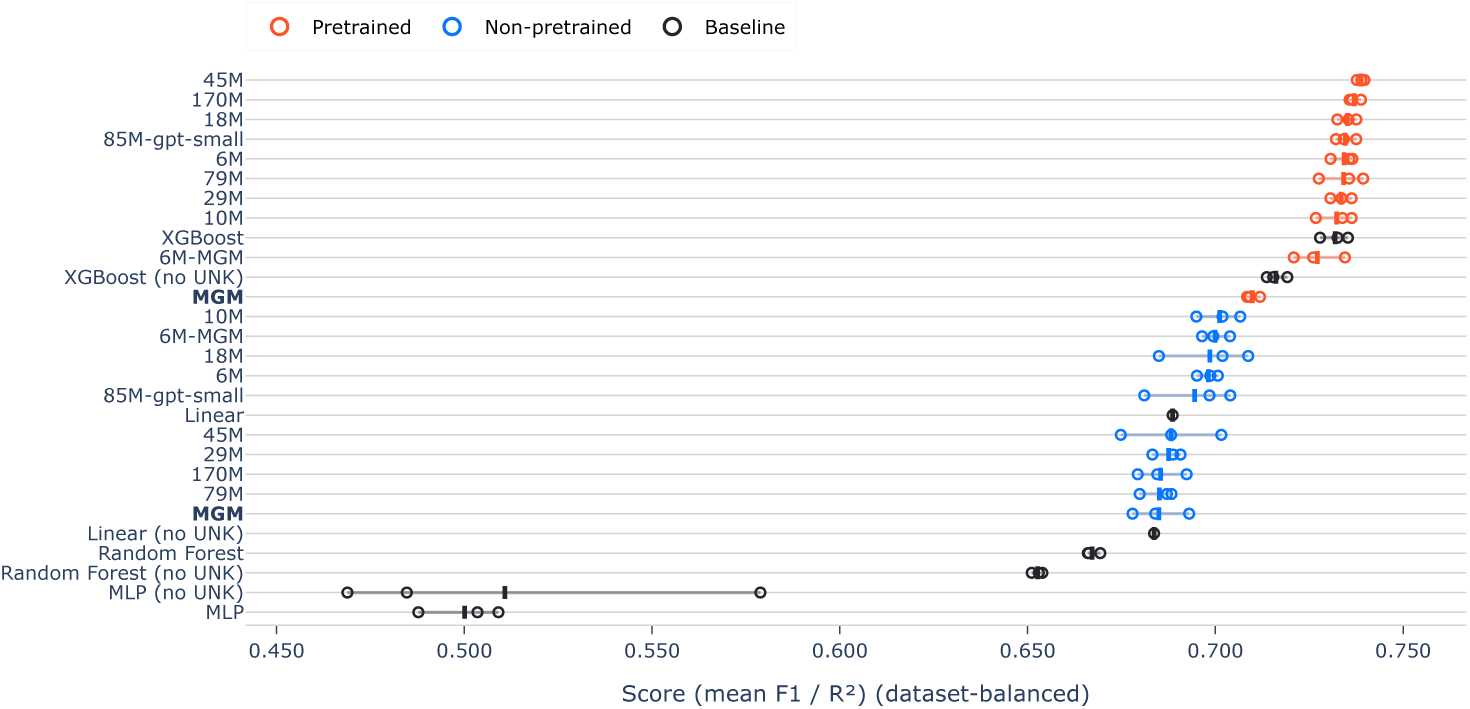
Overall Compass scores for all models, taken as the dataset balanced average of the scores across the 8 benchmark tasks, where classification tasks (1-5 and 7-8) are scored using macro averaged F1 and the regression task (6) is scored using *R*^2^. Solid lines indicate mean score and the hollow circles individual replicates. Pretrained models consistently outperform non-pretrained models and the baselines. Because the benchmarking datasets contain taxonomic labels not seen during pretraining, some tokens are unknown to the transformer models. The baseline methods can use all tokens; when marked as (no UNK), the unknown tokens are removed to match the transformer models’ vocabulary and enable direct comparison.

The per-task breakdown (Figure 3) reveals that the benefit of pretraining is consistent across all tasks. To further illustrate the effectiveness of pretraining for every model we plot the difference in mean score between the pretrained and non-trained models for all tasks (Figure S3), for almost all models across almost all tasks we see that the performance of the pretrained model is superior. This demonstrates the effectiveness of self supervised pretraining to learn general representations of microbiomes that improve performance across a wide variety of downstream tasks. From the per task breakdown we also see that the "no UNK" variant of the XGBoost clearly out-performs all transformers for task 4 and task 7, demonstrating that for some of the small per task datasets we see no clear advantage for using transformers over baseline models even when vocabulary is kept consistent. Supplementary figures provide additional complementary metrics for the classification tasks.

**Figure 3:**
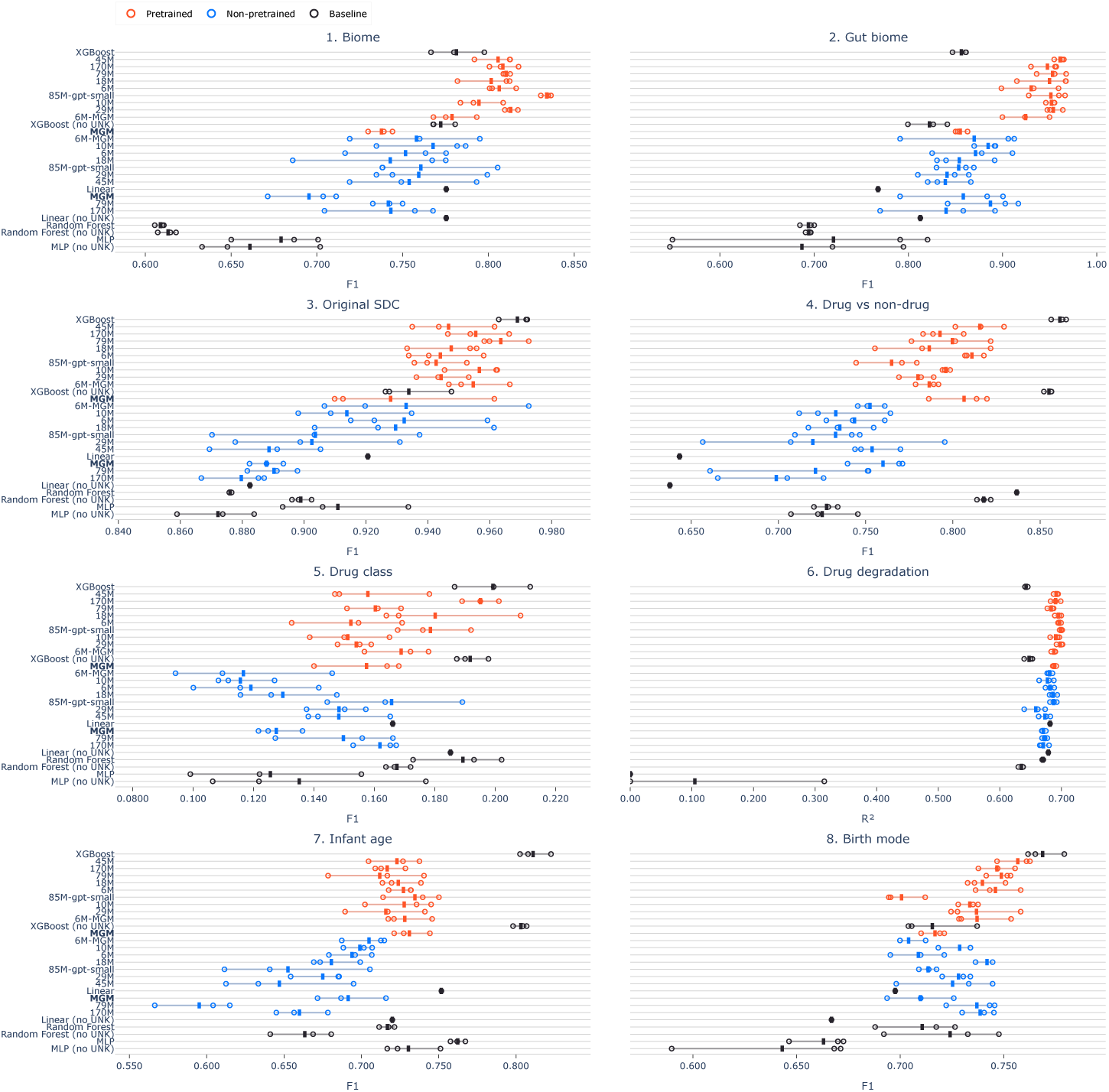
Score per task for all models, classification tasks (1-5 and 7-8) are scored using macro averaged F1 score and the regression task (7) is scored using *R*^2^. Solid lines indicate mean score and the hollow circles individual replicates. Pretrained models consistently outperform non-pretrained models, but for some tasks some baselines outperform the transformer models. Because the benchmarking datasets contain taxonomic labels not seen during pretraining, some tokens are unknown to the transformer models. The baseline methods can use all tokens; when marked as (no unk), the unknown tokens are removed to match the transformer models’ vocabulary and enable direct comparison.

One-vs-one macro ROC AUC (Figure S4), macro precision–recall (Figure S5) and balanced accuracy (Figure S6).

### 2.6 Pretrained transformer models outperform baselines at obtainable dataset sizes

To understand how model performance scales with the amount of available task specific training data, we plot the difference in the benchmark score of the best performing Waypoint model (45M) compared to the XGBoost baselines against benchmark dataset size. Figure 4 shows the difference in mean score (averaged over the three independent runs) between Waypoint and XGBoost as a function of training set size for each task for both the no UNK (Figure 4A) and standard (Figure 4B) XGBoost variants. We see that Waypoint relative performance increases as the number of training examples grows. Notably, from approximately 10k training examples onwards, our Waypoint model consistently achieves higher mean scores than both baselines. Importantly, 10k examples falls within practically obtainable dataset sizes for microbiome applications. The pretrained model outperforms the non-pretrained model across the full range of dataset sizes examined, suggesting that pretraining provides a stable benefit within the range of dataset sizes in the benchmark. Notably, at small dataset sizes (below 1k examples), the Waypoint model underperforms even the no UNK baseline, highlighting that a minimum quantity of training data is required before transformer-based approaches become competitive. These results suggest that Waypoint models are a practical choice for the tasks studied here whenever training datasets of at least 10k examples can be assembled.

**Figure 4:**
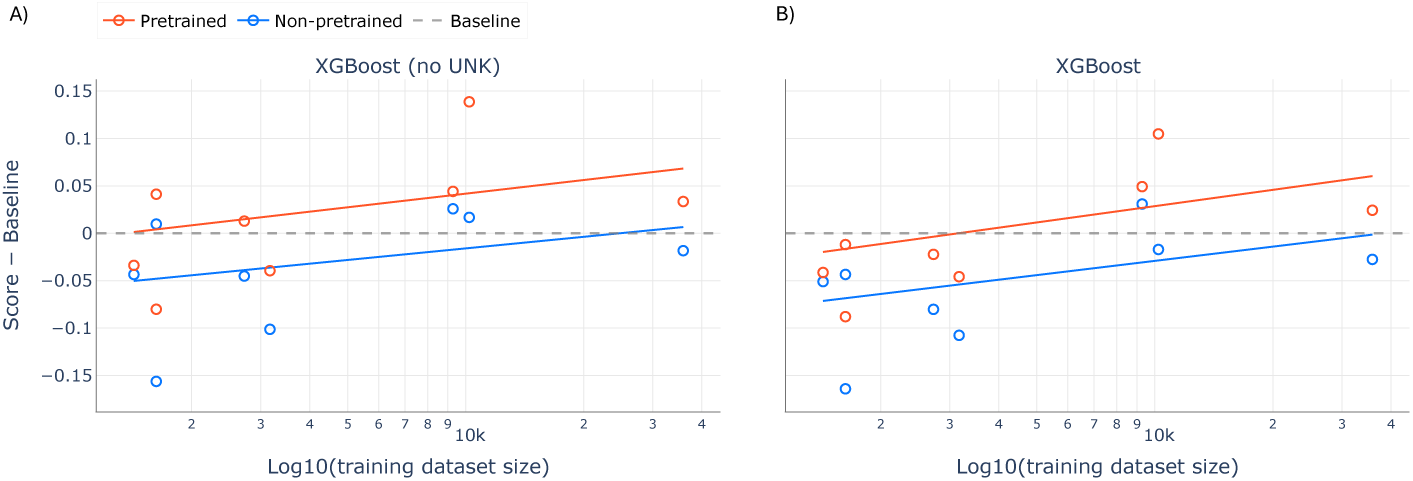
The difference in mean score of transformer models compared to the mean score of XGBoost (no UNK) (A) and XGBoost (B) baselines, against the size of the training dataset for each task in the benchmark. Because the benchmarking datasets contain taxonomic labels not seen during pretraining, some tokens are unknown to the transformer models and here we plot both the XGBoost model with these tokens removed (the no UNK variant) and the standard XGBoost variant with all tokens. At 10k datapoints and above, all pretrained transformers are better than both baselines. Solid lines are ordinary least squares lines of best fit.

### 2.7 Pretraining enables model scaling

We next investigate how benchmark performance varies as a function of model size, and in particular whether pretraining alters scaling behavior. Figure 5 shows benchmark scores for the three replicates for both the pretrained and non-pretrained transformer models. A clear divergence emerges between pretrained and non-pretrained models as model size increases. Pretrained models exhibit a modest but consistent improvement in performance with increasing parameter count, as indicated by the positive slope of the fitted trend. In contrast, non-pretrained models show a degradation in performance as model size grows, with larger models failing to translate additional capacity into improved task performance. This difference suggests that pretraining is critical for effectively utilizing increased model capacity. Without pretraining, larger models appear harder to optimize and may overfit or fail to generalize, leading to negative scaling with increased size. Conversely, pretraining provides a strong initialization that allows larger models to leverage their additional representational capacity, resulting in steady performance gains. This finding has practical implications for model selection: investing in larger architectures is only justified when pretraining is also employed, as scaling a randomly initialised model may even be detrimental. Scaling results for each task are shown in S7, which shows that performance as a function of model size and the effect of pretraining can be variable between tasks, even for those derived from the same underlying datasets.

**Figure 5:**
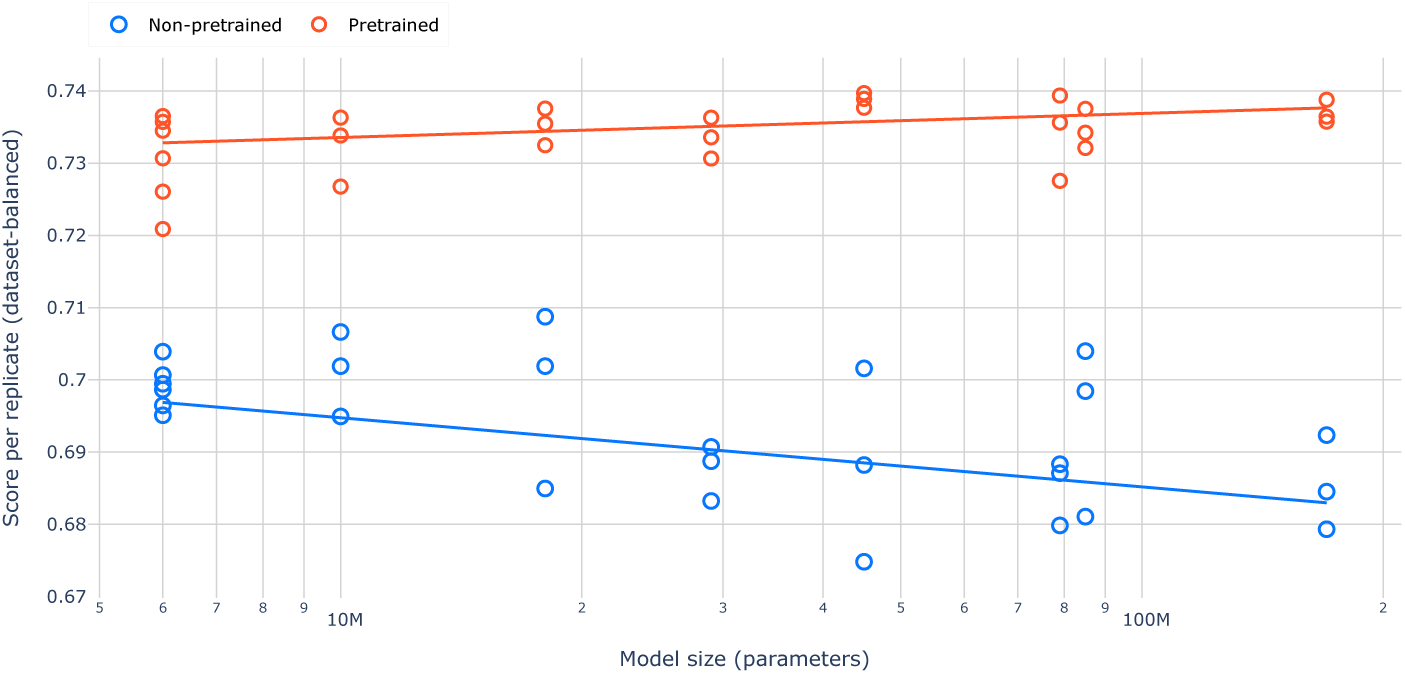
Benchmark performance as a function of model size for pretrained and non-pretrained transformer models. Each point represents the score from an individual training replicate, solid lines are ordinary least squares lines of best fit. Pretrained models show consistent improvements with increasing parameter count, while non-pretrained models exhibit a decline in performance at larger scales, highlighting the importance of pretraining for effective model scaling.

### 2.8 Pretraining generalises between microbiome domains

The Atlas corpus spans a broad range of ecological contexts, from host-associated to marine, freshwater, terrestrial, and engineered environments (Figure 6A). The Compass benchmark, in contrast, is dominated by gut and other host-associated tasks. This raises a natural question: do the gains from pretraining derive specifically from in-domain (gut or human) samples, or do learned representations transfer across biomes? To answer this, we pretrained the 6M Waypoint model on five subsets of Atlas, defined from the biome lineage metadata (Figure 6; see Methods): the complete corpus (*Full-pretrained*); only human-associated samples (*Only human*); only gut samples (*Only gut*); all samples except human-associated (*Exclude human*); and all samples except gut (*Exclude gut*). Any datapoints with missing labels such that their membership in the set was ambiguous were removed to prevent unintentional contamination of the data. Every subset model used the identical architecture, tokeniser, and pretraining and fine-tuning procedure as the standard 6M model, and each was evaluated on the full Compass benchmark across three replicates. Because the subsets have varying numbers of data points and necessarily contain less data than the full dataset (Table 5) we also pretrained five additional models in which each dataset was randomly subsampled to 110k datapoints to control for the effect of dataset size on model performance. For all subset models 90% of the corresponding pretraining data subset was used for training and the remaining 10% for validation. Additionally, we plotted the aggregate benchmark score excluding task 1, this is because task 1 is the only task in the benchmark which is not focused on the human gut microbiome and so removing this allows us to more clearly asses the effect of different pretraining datasets within this domain.

**Figure 6:**
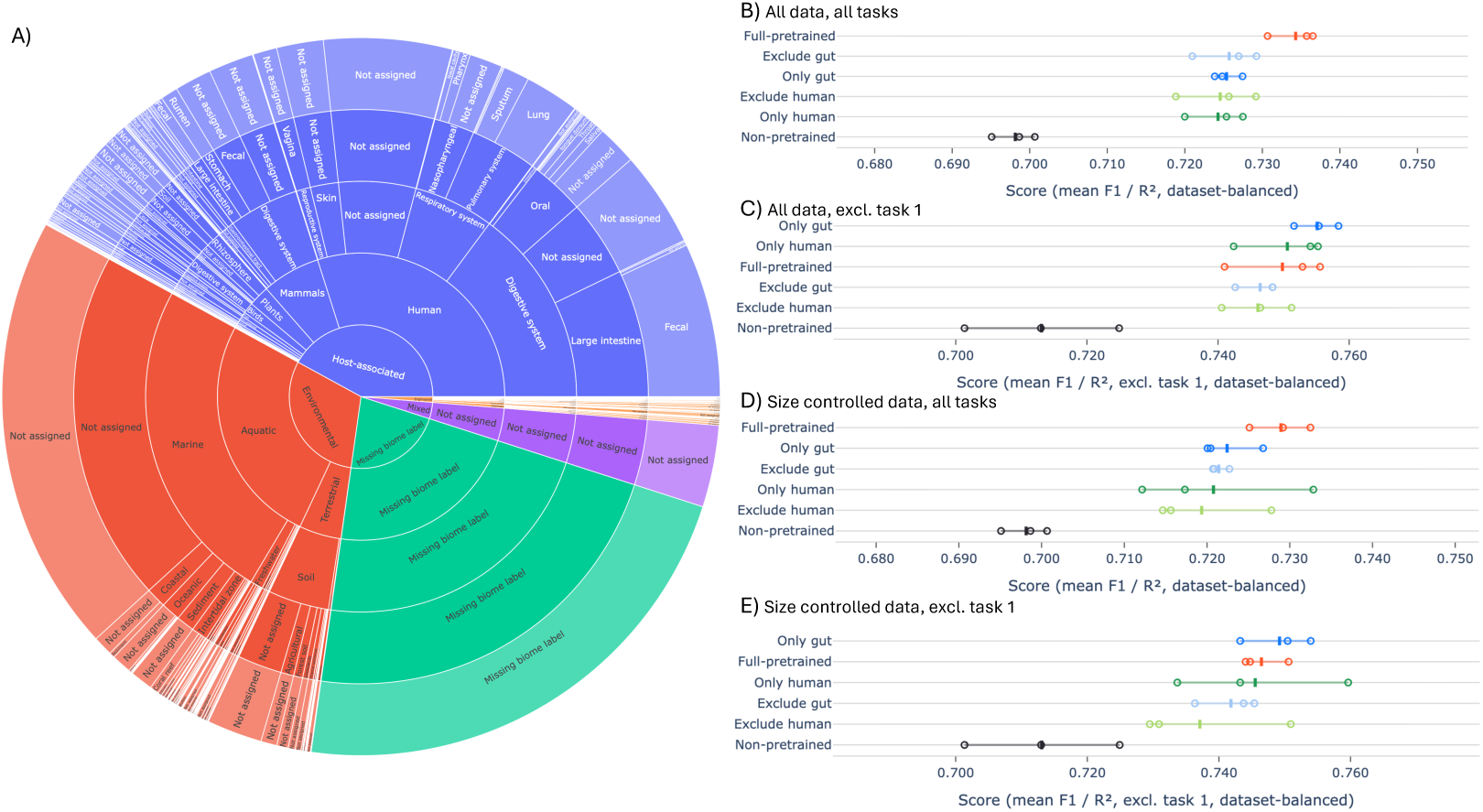
A) Composition of the Atlas pretraining corpus by biome lineage. Each ring corresponds to a level of the MGnify biome ontology. Segment size is proportional to the number of samples. B,C,D,E) Compass scores for models Waypoint-6m models pre-trained on different datasets and evaluated on different tasks.

**Table 5:** Pretraining subsets of Atlas used in the cross-biome transfer experiments. The *human* domain comprises all `Host-associated: Human` samples and the *gut* domain as any data point where the third biome level is equal to Digestive system, these can be both human and non human.

| Subset | Pretraining samples |
| --- | --- |
| Full-pretrained | 539,308 |
| Only human | 154,094 |
| Only gut | 112,709 |
| Exclude human | 246,511 |
| Exclude gut | 287,896 |
| Missing biome metadata | 138,703 |

The mean benchmark score for each pretraining dataset without sub-sampling is shown in Figure 6B. Every pretrained condition substantially outperforms the non-pretrained model, with the model trained on the full corpus performing the best. Strikingly, the four subset models cluster tightly together and are essentially indistinguishable from one another. The greater performance of the *Exclude gut* and *Exclude human* models than the non-pretrained model demonstrates that transfer learning between microbiome domains is effective, indicating that the representations learned during autoregressive pretraining are largely biome-agnostic and transfer across ecological contexts, rather than encoding narrow, domain-specific structure. This advantage is unambiguous: in every panel of Figure 6 the *Exclude* models significantly outperform the non-pretrained model. Removing task 1 from the benchmark (Figure 6 C) notably, but expectedly, lowers the relative scores of *Full-pretrained* and *Exclude gut* such that the gut and human focused datasets are now at the top, however we still see clear transfer learning in comparison to the non-pretrained model. The results from the size controlled pretraining datasets (Figure 6 D, E) show very similar trends, with the expected decrease in score across all pretrained conditions due to lower dataset sizes (the non-pretrained model is unchanged as it uses no pretraining data). The per-task breakdowns (Figure S8, Figure S9) confirms that this pattern is consistent across the benchmark. For most tasks, all pretrained subsets outperform the non-pretrained model, and the in-domain-only and out-of-domain (*Exclude*) subsets overlap heavily, with differences between subsets generally falling within the replicate-to-replicate variation.

Taken together, the results in this section demonstrate that pretraining transfers across ecological contexts: models that never saw gut samples, or never saw human-associated samples, still recover the large majority of the benefit of pretraining on a benchmark dominated by human gut tasks, outperforming the non-pretrained model. The representations a Waypoint model learns from a microbial community remain useful well outside the ecological context it was trained on. This is an encouraging result for microbiome modelling in general, where public data is abundant for a handful of well-studied environments and sparse for most others: it suggests that models for under-sampled biomes need not wait for domain-specific data at scale, and can instead inherit most of their representational quality from whatever broad corpus is available.

## 3 Discussion

In this work we introduced Atlas, Compass and the family of Waypoint models. Respectively, these represent the largest microbiome pretraining dataset assembled to date, a curated benchmark of eight predictive tasks spanning gut and environmental microbiome applications, and the current state of the art in microbiome foundation models. We systematically evaluate the performance of pretrained and non-pretrained GPT-2-style transformer models across a range of model and dataset sizes. Our results demonstrate four principal findings: that self-supervised pretraining on large-scale microbiome data consistently improves downstream task performance, that pretrained transformers outperform classical baselines at practically obtainable dataset sizes, that pretraining is a prerequisite for positive model scaling and that pretrained representations generalise between microbiome domains. These results conclusively demonstrate that autoregressive pretraining on taxonomic abundance sequences is an effective strategy for learning general microbiome representations. Across all eight benchmark tasks, pretrained transformers consistently outperform non-pretrained counterparts. Importantly, our 6M-MGM model, which shares the same architecture as the original MGM model [10], but was pretrained on our larger corpus with fallback tokenisation outperformed the original MGM on the benchmark. This demonstrates that dataset scale and tokenisation strategy are important drivers of foundation model quality.

A practical concern for microbiome foundation models is whether the dataset sizes required to outperform classical baselines are realistically achievable in typical research settings. Our analysis demonstrates that pretrained Waypoint models exceed the XGBoost baseline from approximately 10k training examples and above, even when subject to the disadvantages due to token mismatch. 10k labelled samples is a threshold that falls within the range of many existing microbiome studies, and is likely to become increasingly attainable with advances in large-scale microbiome cohort studies and in vitro microbiome screening. Notably, the dataset exploring how diverse microbiomes metabolize drugs, characterised by [19] (used for task 6 in our benchmark), contained 15419 samples, with 9282 used for training. This finding provides practical guidance for practitioners deciding whether to adopt transformer-based models for a given task.

A central finding of this study is the importance of pretraining to unlock the benefits of model scaling. Without pretraining, increasing model size leads to degraded performance and higher variance across replicates. In contrast, pretrained models exhibit positive scaling behaviour, with larger models achieving better or comparable performance. This highlights that pretraining is necessary for effectively utilising increased model capacity in microbiome settings, meaning that investment in larger transformer architectures for microbiome tasks is only warranted when a sufficiently large pretraining corpus is available. The scaling behaviour observed among pretrained models, with the 45M parameter model achieving the highest mean benchmark score rather than the largest 170M model, warrants further investigation. One possible explanation is that the 170M model requires more task-specific data than is available in the benchmark tasks to fully utilise its capacity, and that its advantage would become apparent at larger dataset sizes. Given that the scores were relatively close, it is also possible that performance has plateaued at this scale, or that the observed differences are due to noise. Future work with larger fine-tuning datasets will be needed to distinguish between these possibilities.

Our cross-biome transfer experiments show that the benefits of pretraining are not confined to the ecological domains represented in the pretraining dataset. Models pretrained without human-associated or gut samples retained most of the downstream benefit of pretraining on a benchmark dominated by human gut tasks and consistently outperformed the non-pretrained model. Moreover, the similar performance of domain-matched and out-of-domain corpora at equal dataset sizes indicates that the learned representations capture broadly transferable properties of microbial communities rather than narrowly domain-specific structure. This further motivates the foundation model paradigm in a field where labelled data is scarce for many biomes: as Atlas and comparable resources grow, models pretrained on them should extend to new and underrepresented environments rather than being limited to the domains best represented in the corpus.

Several limitations of this study should be acknowledged. First, Compass currently focuses exclusively on gut and environmental microbiome tasks; while the pretraining corpus is ecologically diverse, the benchmark itself does not yet reflect the full breadth of microbiome applications, including oral, skin, or respiratory microbiomes. Expanding the benchmark to cover a wider range of body sites and environments is an important direction for future work. Second, we were unable to include comparisons with two recently published microbiome foundation models [11, 12] due to the absence of publicly available models, which limits the scope of our comparative analysis. We encourage the release of model weights alongside future publications to facilitate community benchmarking. Furthermore, our models currently have two obvious limitations which should be addressed in subsequent work; they are not aware of relative abundance values beyond the rank ordering of taxa and they are vulnerable to token mismatch between the pretraining and finetuning datasets. Previous papers have suggested the efficacy of more fine-grained treatment of relative abundance [12], although other work has suggested that rank ordering is just as effective [11]. We see that token mismatch is an issue of varying magnitude with all of our benchmarking datasets (Table 4), likely limiting the efficacy of our models. This can be solved by reprocessing raw data with consistent bioinformatics pipelines. However, this can require significant expertise and compute, so future work to generalise tokenisation schemes to work across datasets should be done. Additional work should also explore a broader range of model architectures beyond GPT-2 to assess their suitability for microbiome modelling tasks.

Taken together, our results establish that self-supervised pretraining on large-scale microbiome sequence data is a powerful and practical strategy for improving performance across diverse downstream tasks and enables foundation models to learn general representations of microbiomes. The release of the Atlas dataset will enable the community to pretrain their own microbiome foundation models. The Compass benchmark provides a standardised and reproducible framework for evaluating future models, and we hope it will serve as a community resource for tracking progress in microbiome machine learning. We have also demonstrated that Waypoint models have achieved state of the art performance and we are releasing the Waypoint-6M, Waypoint-45M and Waypoint-170M foundation models to the community. As microbiome datasets continue to grow in scale and diversity, we anticipate that the benefits of foundation model pretraining will become increasingly pronounced, making this a productive direction for future research.

## 4 Methods

### 4.1 Pretraining Dataset

#### Data Acquisition

Taxonomic abundance data were downloaded from the MGnify public database [16] across all available pipeline versions (v1.0–v5.0) and four sequencing modalities: amplicon 16S rRNA, whole-genome shotgun metagenomic, metagenomic assembly, and metatranscriptomic. Data were obtained in the form of per-study abundance matrices where each column is a sample run accession and each row is a taxon, with values representing relative abundances. All datasets were versioned with DVC to ensure reproducibility of all download and processing steps.

Two quality filters were applied: (i) any taxon with relative abundance below 10*^−^*^4^ was removed, and (ii) any sample with fewer than 10 taxa remaining after the abundance filter was removed. After filtering, the dataset comprised 539,308 datapoints.

Prior to any tokenisation or model training, the filtered dataset was randomly partitioned into a pretraining set (90%) and a held-out bench-marking set (10%) using a fixed random seed (*s* = 42). The benchmarking partition is used exclusively to supply labelled samples for tasks 1 and 2 of the benchmark, ensuring no overlap with pretraining data.

#### Taxonomic Tokenisation

Each microbiome sample is represented as an ordered sequence of taxonomic strings. A custom `TaxonomicTokenizer` was built by extracting all genus-level names present in the pretraining data. Where a taxon lacked a genus-level classification, the tokeniser falls back to the most specific available rank (family, order, and so on). The vocabulary comprises 14,389 taxonomic tokens plus five special tokens (`<bos>, <eos>, <pad>, <unk>, <mask>`), for a total vocabulary size of 14,394.

As in [10], taxa within each sample are ordered by their z-scored relative abundances, calculated using statistics from the pretraining corpus. For taxon (t) in sample (i), the z-score is calculated as 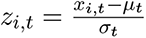, where (*x_i,t_*) is the observed relative abundance of taxon (t) in sample (i), and (*µ_t_*) and (*σ_t_*) are its mean and standard deviation, respectively, across the pretraining corpus. Tokens are then sorted from highest to lowest z-score. The means and standard deviations of each token in the pretraining dataset are serialised with the tokeniser, allowing the same normalisation to be applied to downstream tasks. Sequences are truncated to a maximum length of 510 tokens and then flanked by `<bos>` and `<eos>` tokens (to give a total maximum length of 512 tokens). The vocabulary is initialised during pretraining using all taxa in the pretraining dataset. During downstream benchmarking, unknown taxa are assigned the `<unk>` token and appended to the sequence after the z-score-sorted known tokens.

### 4.2 Pretraining

We pretrain GPT-2-style causal language models with a next-token prediction objective. We explore model scales from 6M to 170M parameters by varying the number of layers and the embedding dimension. All models use HuggingFace `transformers` and were trained using the AdamW optimiser with a linear learning-rate warmup over the first 1,000 steps (learning rate 10*^−^*^3^, weight decay 10*^−^*^3^, batch size 32 per device). Training proceeded for up to 100 epochs with early stopping on validation loss (patience of 10 evaluations at half-epoch intervals). A 10% random split of the pretraining set was held out as an in-training validation set for early stopping; this split is distinct from the held-out benchmarking partition. The GPU used and runtime of each pretraining run are shown in Table 6.

**Table 6:**
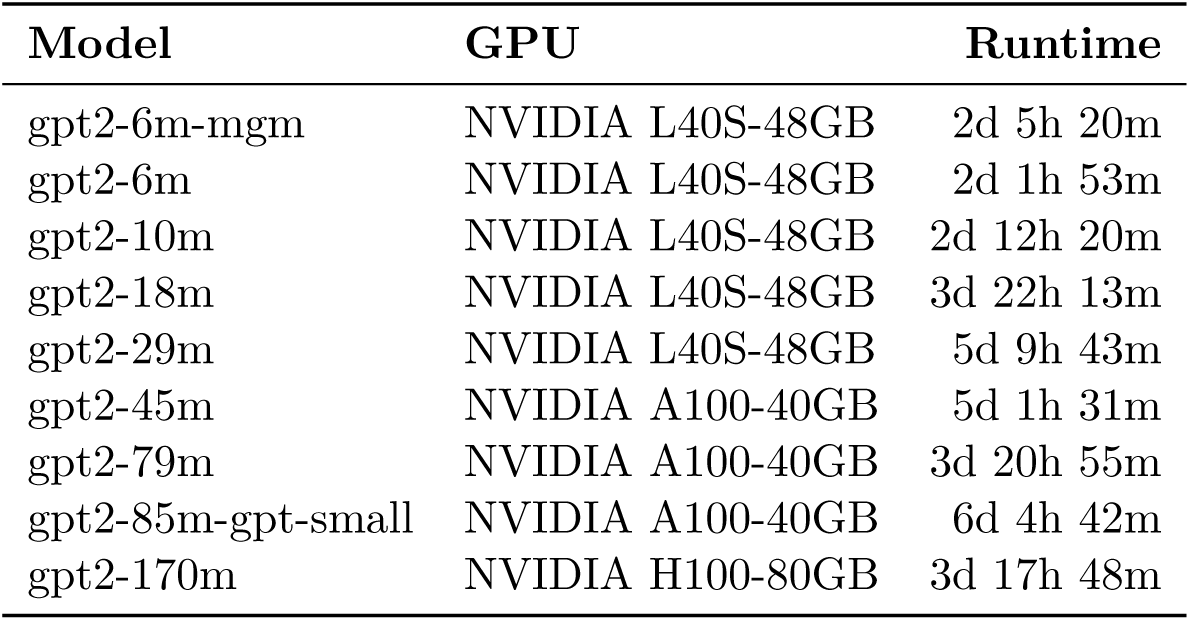
GPU and runtime for model pretraining.

| Model | GPU | Runtime |
| --- | --- | --- |
| gpt2-6m-mgm | NVIDIA L40S-48GB | 2d 5h 20m |
| gpt2-6m | NVIDIA L40S-48GB | 2d 1h 53m |
| gpt2-10m | NVIDIA L40S-48GB | 2d 12h 20m |
| gpt2-18m | NVIDIA L40S-48GB | 3d 22h 13m |
| gpt2-29m | NVIDIA L40S-48GB | 5d 9h 43m |
| gpt2-45m | NVIDIA A100-40GB | 5d 1h 31m |
| gpt2-79m | NVIDIA A100-40GB | 3d 20h 55m |
| gpt2-85m-gpt-small | NVIDIA A100-40GB | 6d 4h 42m |
| gpt2-170m | NVIDIA H100-80GB | 3d 17h 48m |

### 4.3 Benchmark Construction

#### Datasets

The benchmark draws on four publicly available microbiome datasets. The MGnify dataset [16] consists of metagenomically profiled samples spanning environments including gut, skin, respiratory tract, oral cavity, marine, freshwater, soil, and engineered systems; biome labels were parsed from the hierarchical biome lineage metadata field (e.g., `root: Host-associated: Human: Digestive system: Fecal`). The Shi et al. drug–microbiome dataset [18] comprises 16S rRNA amplicon profiles of stabilised gut microbial communities exposed to a panel of drugs or control conditions, with drug identity and ATC classification provided as metadata. The Mastrorilli et al. dataset [19] contains paired community composition and drug degradation rate measurements for human gut microbial communities incubated with a panel of compounds. The Roswall et al. infant dataset [20] contains longitudinal 16S rRNA profiles of infant gut communities with associated delivery mode and timepoint metadata.

#### Data processing

All datasets are processed to a common long format in which each row represents one datapoint, with two list fields storing taxonomic identifiers and corresponding relative abundances in descending order. Taxonomic strings are encoded with standard rank prefixes (e.g., `k Bacteria; p Firmicutes; g Lactobacillus`). ASV-level abundance tables were collapsed to genus level by summing relative abundances within each unique genus-level lineage per sample. For the Mastrorilli et al. dataset, single-rank taxon identifiers were first mapped to full NCBI-derived taxonomic strings prior to genus-level collapse.

#### Train/validation/test splits

Data are partitioned into train (80%), validation (10%), and test (10%) sets using a fixed random seed (42). Because the Mastrorilli dataset is *n_microbiomes_ n_drugs_* each microbiome appears many times in the dataset. To prevent the same microbiome appearing in train and testing data all observations for a given microbiome are grouped before splitting. The Mastrorilli et al. dataset uses a 60/20/20 split. All splits are pre-computed and stored with the processed data to ensure reproducibility across model evaluations.

#### Task definitions

Eight tasks are defined over the four datasets. Tasks 1 and 2 are multi-output classification tasks over the MGnify dataset [16]; task 1 predicts all five biome hierarchy levels simultaneously, while task 2 restricts to samples labeled as *Digestive system* at hierarchy level 3 and predicts only the two finer-grained levels. For datapoints with only partial coverage of the biome levels, the levels that are missing are masked so that they do not contribute to the loss. Tasks 3–5 are single-output classification tasks over the Shi et al. dataset: task 3 predicts the sample from which a drug perturbed community originated from, task 4 predicts binary drug/control status, and task 5 predicts the first-level ATC drug class a microbiome was exposed to. Task 6 is a regression task predicting the rate of drug degradation from microbiome composition and a one-hot-encoded drug indicator. Tasks 7 and 8 are single-output classification tasks over the infant dataset, predicting the age of the infant the microbiome was taken from and the delivery mode respectively.

#### Evaluation metrics

We report macro F1 as the primary summary of performance for classification tasks. This metric computes the F1 score independently for each class and then averages them with equal weight, ensuring that the metric is not over sensitive to performance on majority labels. Supplementary figures report additional complementary metrics; one-vs-one macro ROC AUC (Figure S4) captures pairwise discriminability by averaging AUC values across all class pairs, macro precision–recall (Figure S5) summarizes the balance between precision and recall across classes without being dominated by label frequency, balanced accuracy (Figure S6) reports the unweighted mean of per-class recall, reflecting how well each class is recovered on average. For the regression task in the benchmark, performance is evaluated using the coefficient of determination (*R*^2^). Because *R*^2^ can take negative values we clamp it to 0 to maintain consistency with other metrics reported on a [0, 1] scale.

#### Benchmark score aggregating

To prevent datasets represented by multiple benchmark tasks from disproportionately influencing aggregate performance, we used dataset-balanced averaging. For each model and replicate, task scores were first averaged within each source dataset and these dataset-level means were then averaged across datasets. Specifically, for dataset (d), containing task set (*T_d_*), we calculated

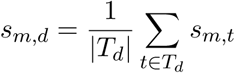

and defined the dataset-balanced benchmark score as

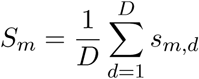

where (*s_m,t_*) is the score of model (*m*) on task (*t*) and (*s_m,d_*) is the score of model (*m*) on dataset (*d*), and (D=4) is the number of source datasets. The four dataset groups comprised tasks 1–2, tasks 3–5, task 6, and tasks 7–8. This gives each source dataset equal weight, irrespective of how many prediction tasks it contributes.

### 4.4 Downstream Fine-tuning on the Benchmark

For each benchmark task, a lightweight task-specific head was appended to the GPT-2 backbone. A sequence-level representation was obtained by taking the hidden state of the last non-padding token (i.e. the <eos> position). For task 6 (drug degradation), which includes drug identity as a feature, a one-hot encoding of the drug compound was concatenated to this before the head. For classification tasks, one independent linear head per target was applied to produce class logits, with cross-entropy loss computed using class-frequency-based weighting to mitigate class imbalance. For the regression task (task 6), a single linear head was used and trained with mean squared error loss.

All model weights (LLM backbone + head) were updated during fine tuning. Models were trained with the AdamW optimiser (learning rate 3 *×* 10*^−^*^5^, weight decay 10*^−^*^3^, batch size 64, linear warmup over 1,000 steps) for up to 300 epochs with early stopping on validation loss (patience of 5 evaluations every 400 steps). For the classification tasks class weights were computed on the training split using the balanced weighting scheme *w_c_* = *N/*(*K n_c_*), where *N* is the total number of training samples, *K* is the number of classes, and *n_c_* is the number of training samples belonging to class *c*. These weights were passed to the cross-entropy loss, weighting classes to counteract class inbalance. The best checkpoint (lowest validation loss) was reloaded before test-set evaluation.

Benchmark samples were tokenised using the same `TaxonomicTokenizer` and z-scored relative abundance ordering described above (Section 4.1), with token z-score statistics loaded from the file saved alongside each model checkpoint during pretraining. This ensures that the token ordering seen at fine-tuning time is consistent with that used during pretraining.

For the MGM model the weights, tokeniser and token z-score statistics were taken from the repository provided in the paper [10].

To benchmark non pretrained version of the models, weights were simply randomly reinitialised prior to benchmarking.

#### Baselines

Four baselines operating on taxon abundance features were included for comparison. All four share a common feature pipeline.

For the "no UNK" variants taxa that the pretrained model cannot represent are first removed, so that the baselines see exactly the same taxonomic information as the tokenizer-based models and the comparison is not confounded by differences in vocabulary coverage. Each taxon name is tokenized with the pretrained model’s tokenizer and dropped if any of its tokens maps to the unknown token; the relative abundances of the surviving taxa in that sample are then renormalised to sum to one.

A bag-of-taxa vector is then constructed for each sample, where each dimension corresponds to a unique taxon and its value is the observed relative abundance (0 if absent). Abundances are then centred-log-ratio (CLR) transformed row-wise with a pseudocount of 10*^−^*^6^, and each taxon dimension is standardised to zero mean and unit variance using training-split statistics. For tasks including drug identity, a one-hot drug encoding is appended after the transform and is itself left untransformed.

For all four models, hyperparameters were selected by grid search on the validation split, scored by cross-entropy for classification and mean squared error for regression, and class imbalance was handled by class-balanced weighting of the loss (*n/*(*K · n_k_*) for class *k*). For logistic regression (classification) and ridge regression (regression), the regularisation strength was selected by grid search over 0.01, 0.1, 1, 10 with a maximum of 1000 solver iterations. For the random forest baseline, the search covered number of trees (100, 200, 300, 500) and maximum depth (10, 20, unlimited), with 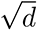 features considered per split. For the XGBoost baseline, maximum tree depth was searched over 4, 6, 8. Because the feature matrix is very wide (approximately 29,000 taxon columns), the number of boosting rounds and the learning rate were fixed to a coupled default of 200 trees at a learning rate of 0.1 rather than scanned, as a lower learning rate simply requires proportionally more trees. Remaining parameters were fixed at row subsampling 0.8, column subsampling per tree 0.2, minimum child weight 1.0, and *L*_2_ regularisation 1.0, using the histogram tree-construction method. The feedforward neural network baseline is a multilayer perceptron with two hidden layers of 256 and 128 units, each followed by a ReLU activation and dropout, and a linear output head. Dropout rate (0.1, 0.3) and learning rate (10*^−^*^3^, 3 10*^−^*^4^) were grid-searched. The network was trained with Adam (weight decay 10*^−^*^5^) at batch size 256 for up to 200 epochs, using weighted cross-entropy for classification and weighted mean squared error for regression. Early stopping used a held-out 10% of the training rows with a patience of 5 epochs, kept separate from the validation split so that the latter is used solely for hyperparameter selection. For multi-target regression a single shared network with a multi-output head was used, whereas the other three baselines fit one independent model per target.

### 4.5 Biome Subset Pretraining

To test whether the benefits of pretraining are specific to in-domain data or transfer across ecological contexts, we pretrained the 6M model on five subsets of the Atlas corpus. Subsets were defined from the hierarchical biome lineage metadata associated with each sample (e.g. `root: Host-associated: Human: Digestive system: Fecal`), the same field used to construct the biome-classification tasks. We defined the *human* domain as all data points whose lineage falls under `Host-associated: Human`, and the *gut* domain as any data point where the third biome level is equal to `Digestive system`, these can be human or non human. The five conditions were: *Full-pretrained*, the complete Atlas pretraining set; *Only human* and *Only gut*, restricted to the respective in-domain samples; and *Exclude human* and *Exclude gut*, comprising all pretraining samples *outside* the respective domain. Any data-points with missing labels such that their membership in a set was ambiguous were automatically removed to prevent any unintentional contamination of the data.

Each subset model was pretrained from scratch using the identical architecture, tokeniser, z-score abundance ordering, optimiser settings, context length, and early-stopping criteria as the standard 6M model (see *Pretraining*). The tokeniser vocabulary was held fixed to that of the full corpus across all conditions so that downstream tokenisation and token coverage are identical, isolating the effect of pretraining-data composition. Each pretrained subset model was then fine-tuned and evaluated on all eight Compass tasks following the same downstream procedure as the main experiments, with three independent replicates per task. The non-pretrained reference model is the randomly-initialised 6M model evaluated under the same fine-tuning protocol.

## Model, Data and Code Availability

The Waypoint model weights, the Atlas dataset and the Compass benchmark are available on Hugging Face at https://huggingface.co/outpost-bio/, and example pre-training and finetuning/benchmarking code is available on GitHub at https://github.com/Outpost-Bio/waypoint. All artifacts are released under the Apache License, Version 2.0. The Hugging Face repositories are gated; access is granted automatically upon request via the Hugging Face interface.

## A Appendix

### A.1 Supplementary Figures

**Figure S1:**
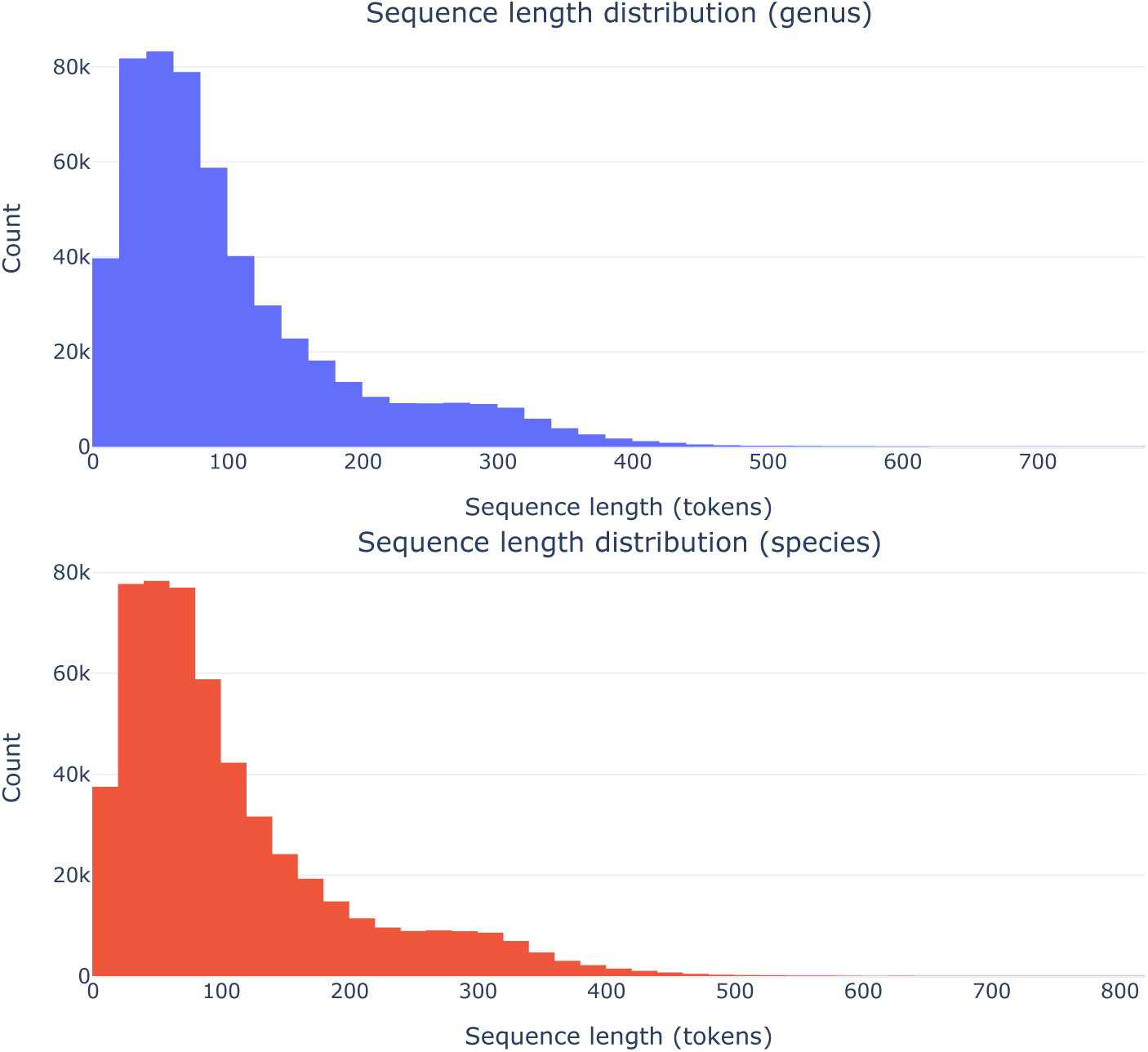
The distribution of sequence lengths when tokenising at the genus (top) and species (bottom) taxonomic levels.

**Figure S2:**
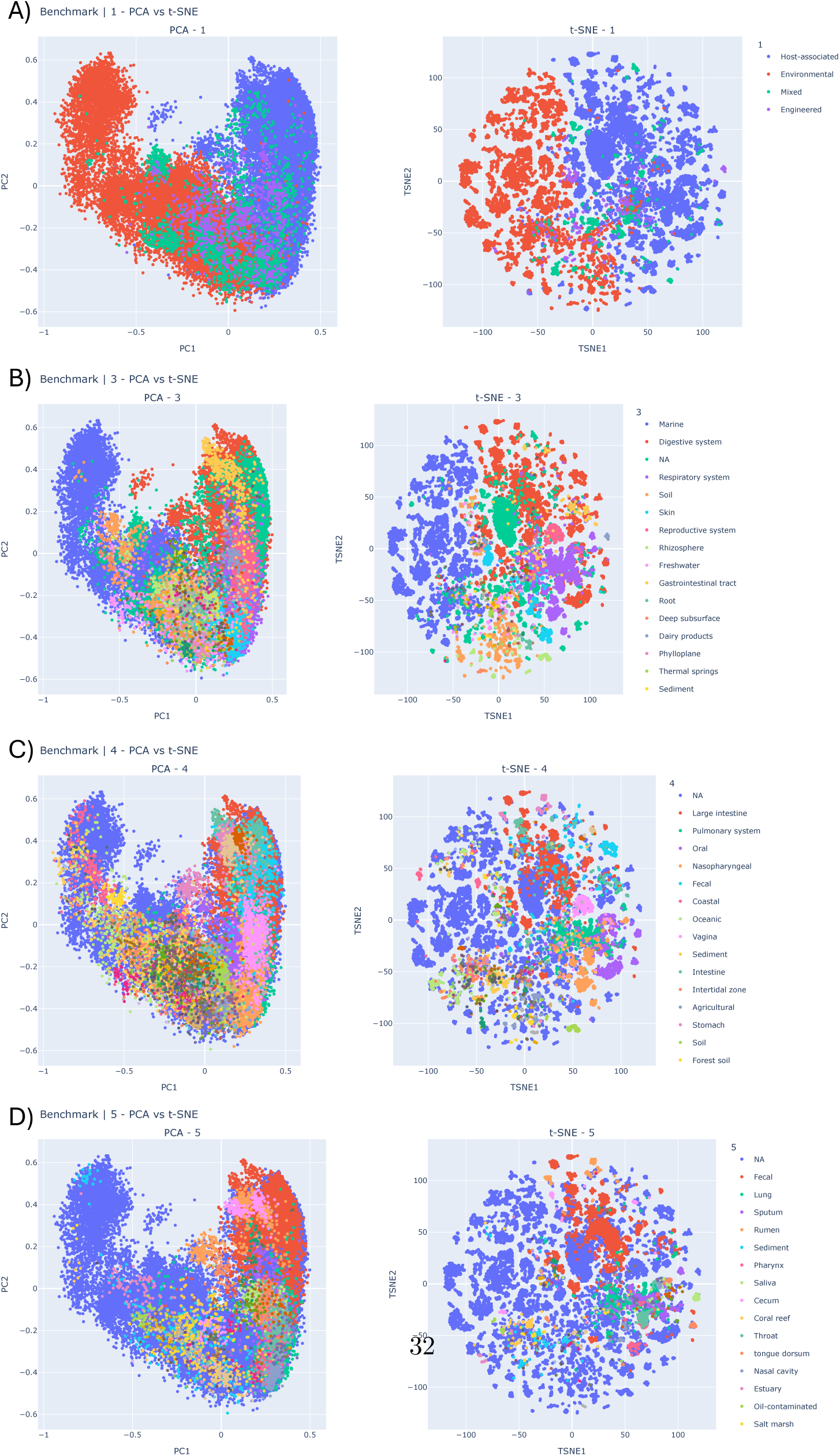
A-D) PCA and t-SNE plots of mean pooled model embeddings coloured according to the 1st, 3rd, 4th, 5th levels of biome hierarchy.

**Figure S3:**
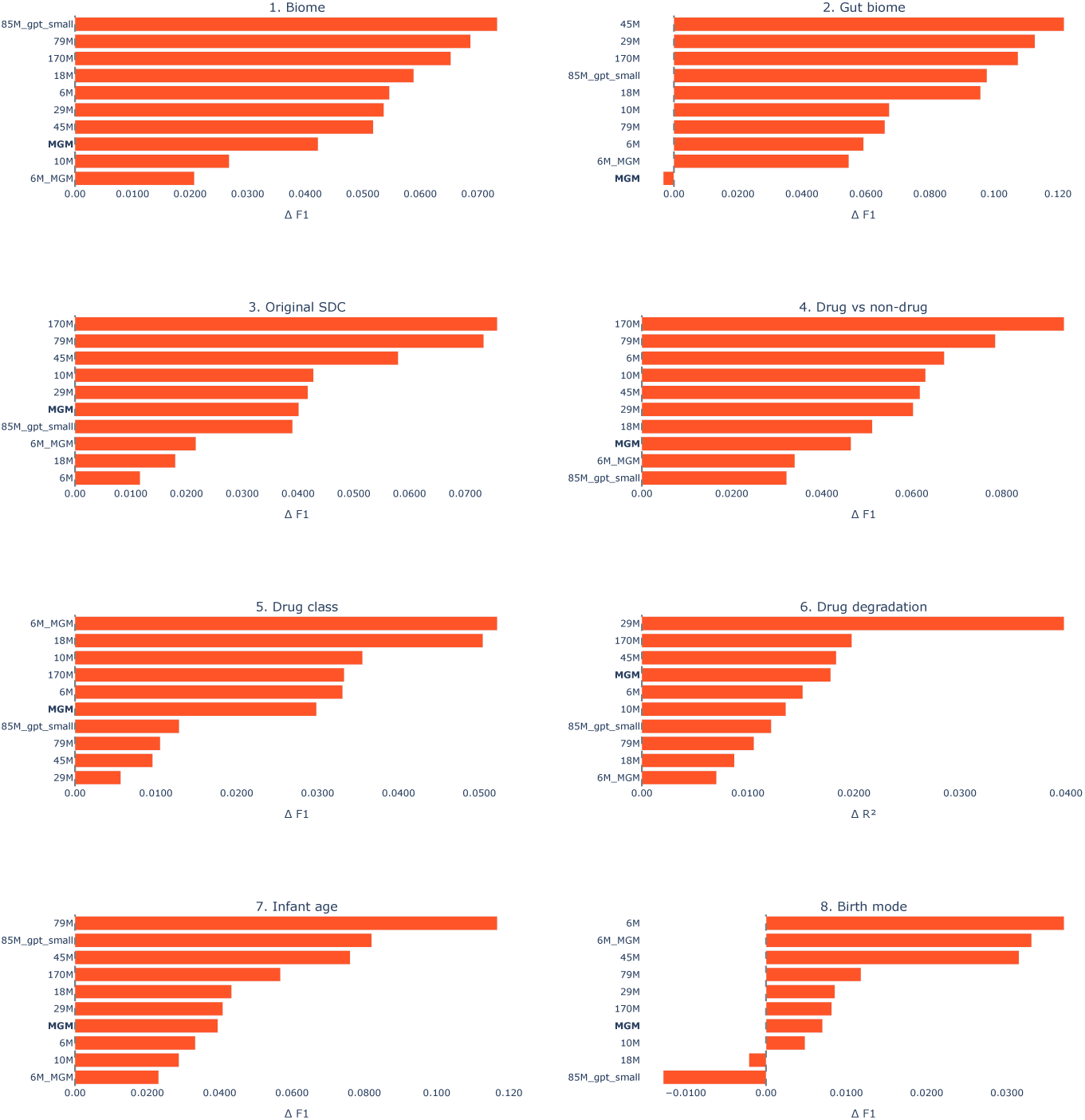
The change in benchmark score caused by pretraining for all tasks and models. Positive values indicate pretrained models perform better, negative values indicate non-pretrained models perform better. For almost all models and all tasks

**Figure S4:**
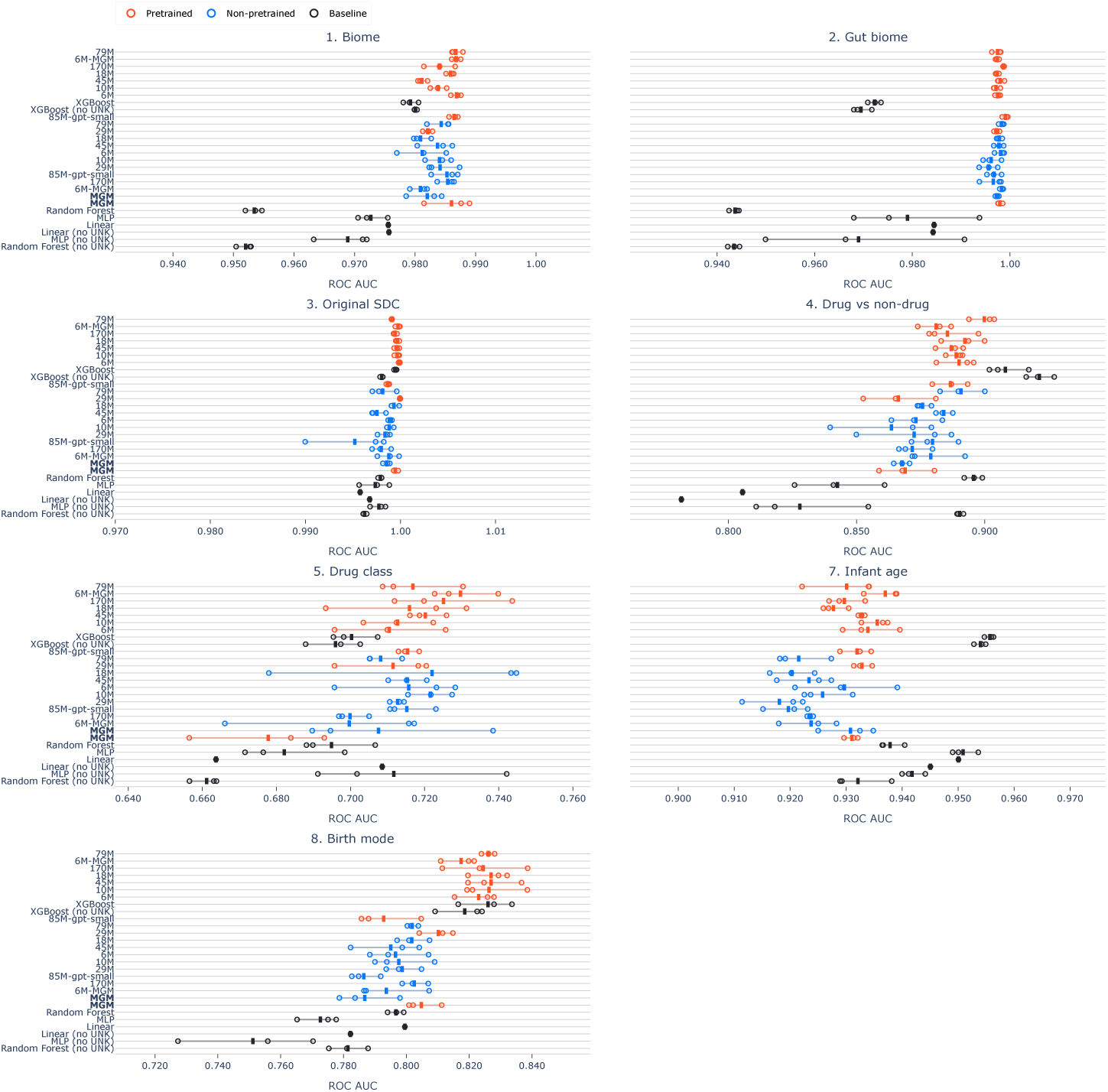
ROC AUC per classification task for all models. Pretrained models consistently outperform non-pretrained models, but we see that for some tasks the RF baseline outperforms the transformer models. Because the benchmarking datasets contain taxonomic labels not seen during pretraining, some tokens are unknown to the transformer models. The baseline methods can use all tokens; when marked as (no unk), the unknown tokens are removed to match the transformer models’ vocabulary and enable direct comparison.

**Figure S5:**
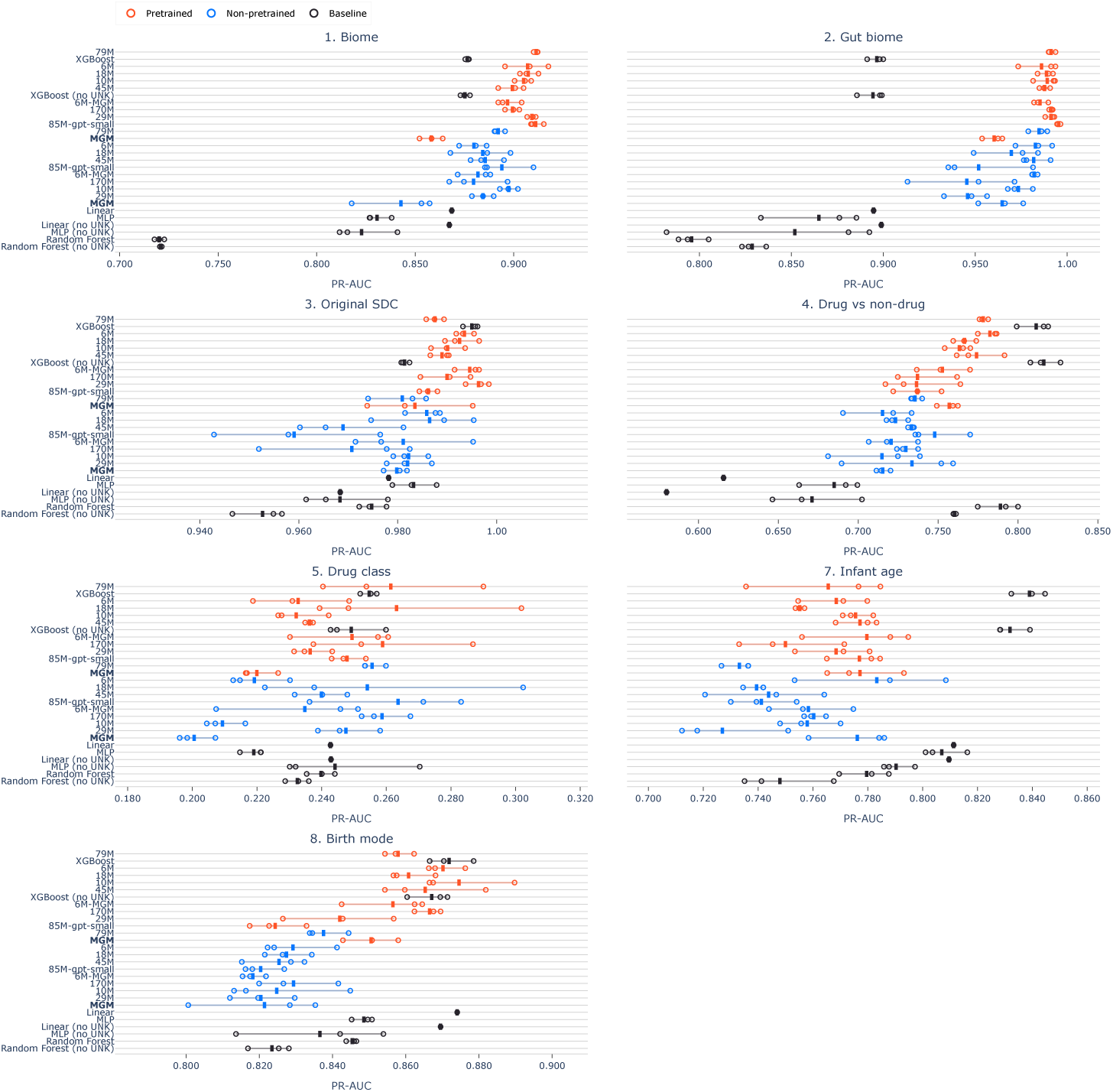
PR AUC per classification task for all models. Pretrained models consistently outperform non-pretrained models, but we see that for some tasks the RF baseline outperforms the transformer models. Because the benchmarking datasets contain taxonomic labels not seen during pretraining, some tokens are unknown to the transformer models. The baseline methods can use all tokens; when marked as (no unk), the unknown tokens are removed to match the transformer models’ vocabulary and enable direct comparison.

**Figure S6:**
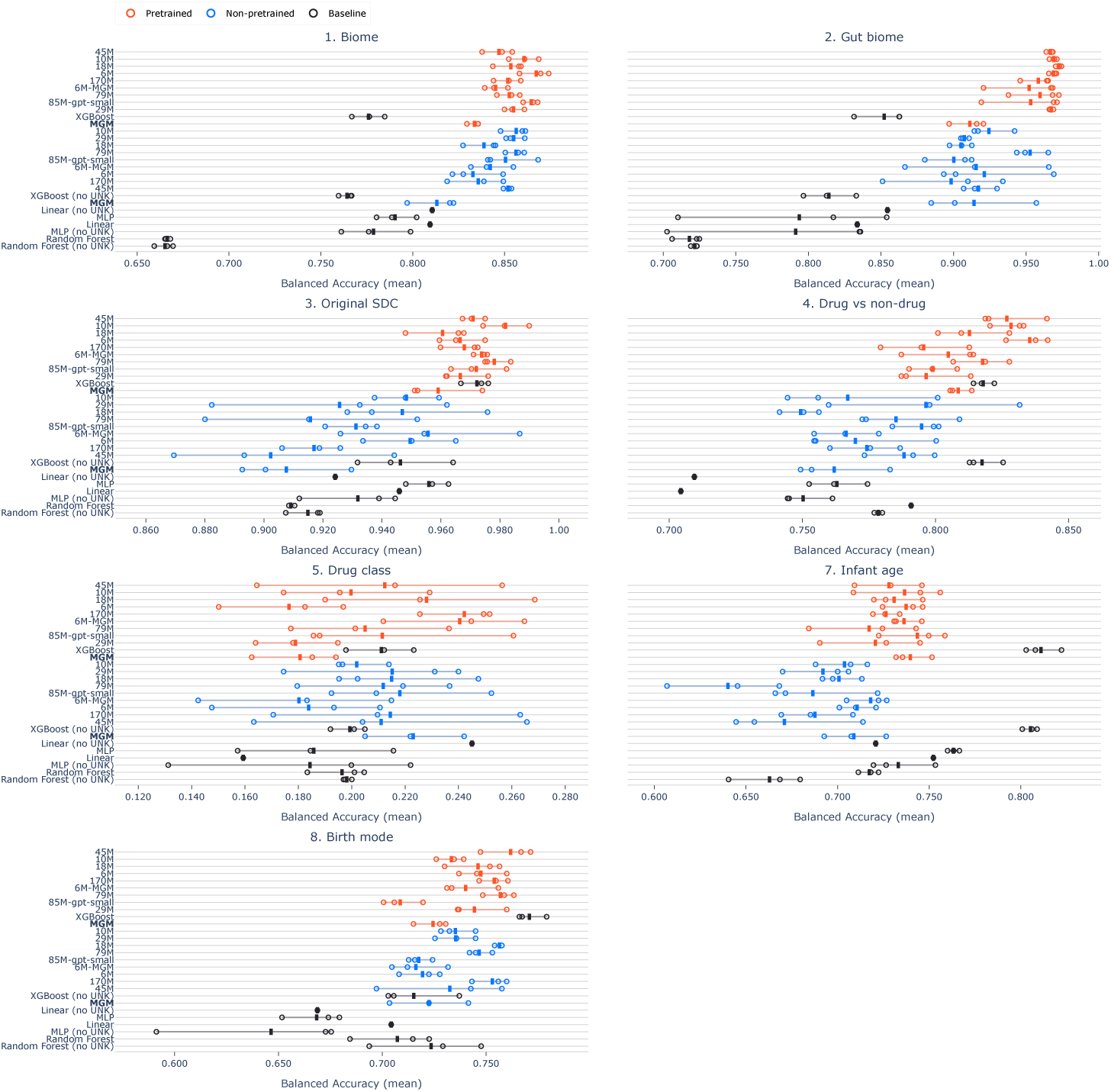
Balanced accuracy per classification task for all models. Pretrained models consistently outperform non-pretrained models, but we see that for some tasks the RF baseline outperforms the transformer models. Because the benchmarking datasets contain taxonomic labels not seen during pretraining, some tokens are unknown to the transformer models. The baseline methods can use all tokens; when marked as (no unk), the unknown tokens are removed to match the transformer models’ vocabulary and enable direct comparison.

**Figure S7:**
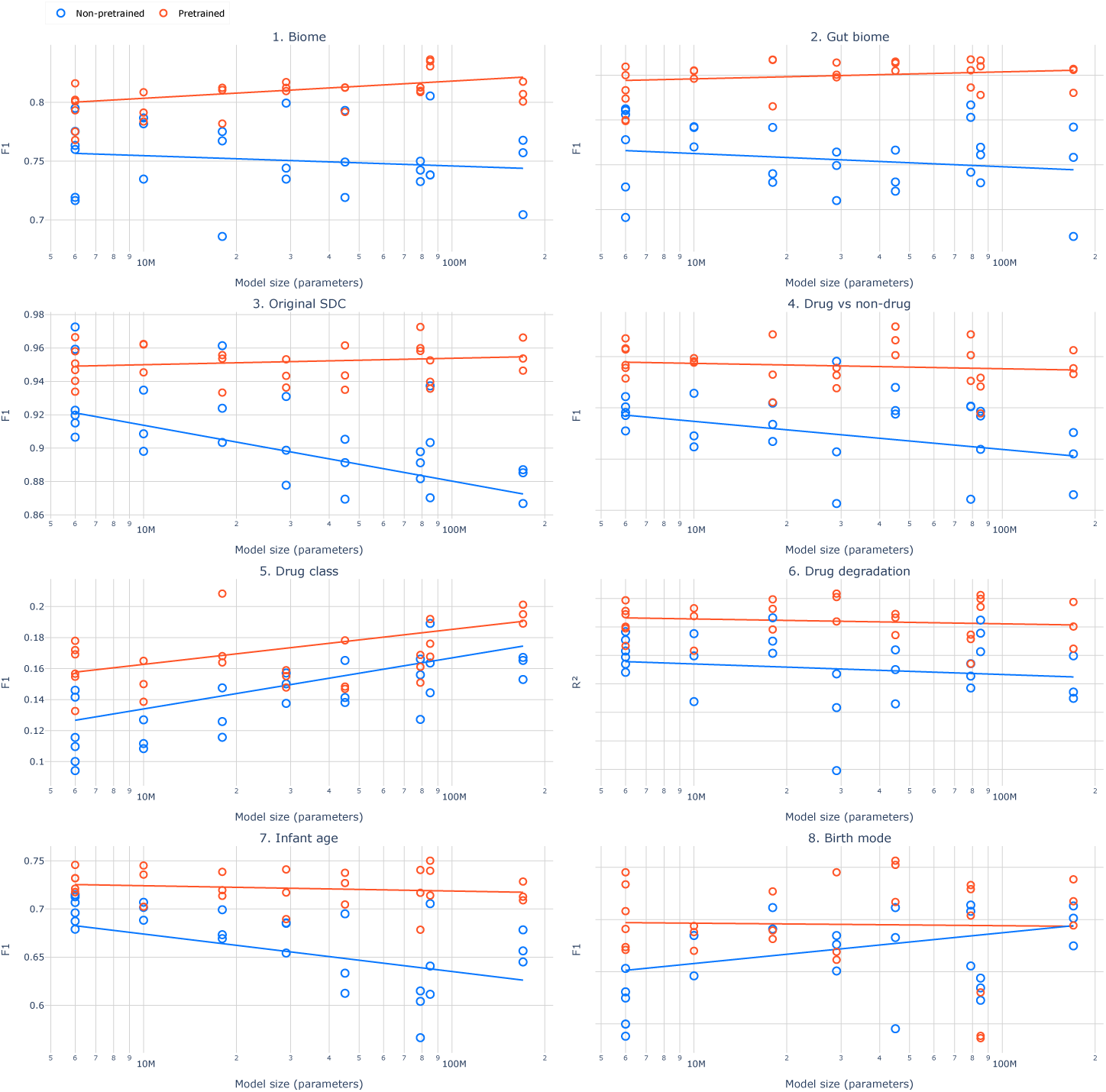
Benchmark performance as a function of model size for pretrained and non-pretrained transformer models for all tasks in the benchmark. Each point represents the score from an individual training replicate, solid lines are ordinary least squares lines of best fit. Scaling behaviour is not uniform across tasks.

**Figure S8:**
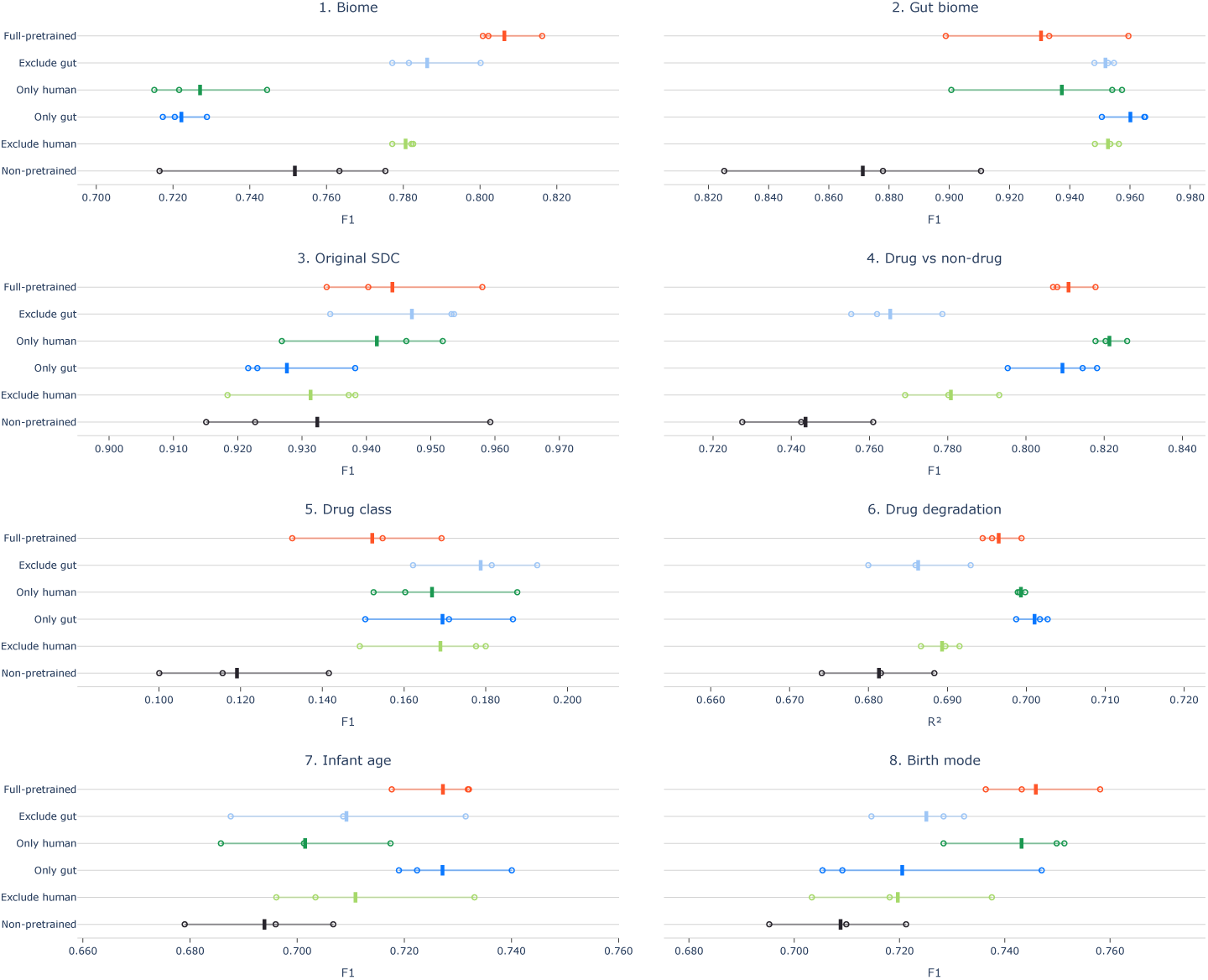
Per-task Compass scores for the 6M model pretrained on different biome subsets of Atlas. Classification tasks (1–5, 7–8) are scored by macro-averaged *F*_1_ and the regression task (6) by *R*^2^. Solid lines indicate the mean and hollow circles individual replicates. The cross-biome transfer observed in the aggregate holds across individual tasks.

**Figure S9:**
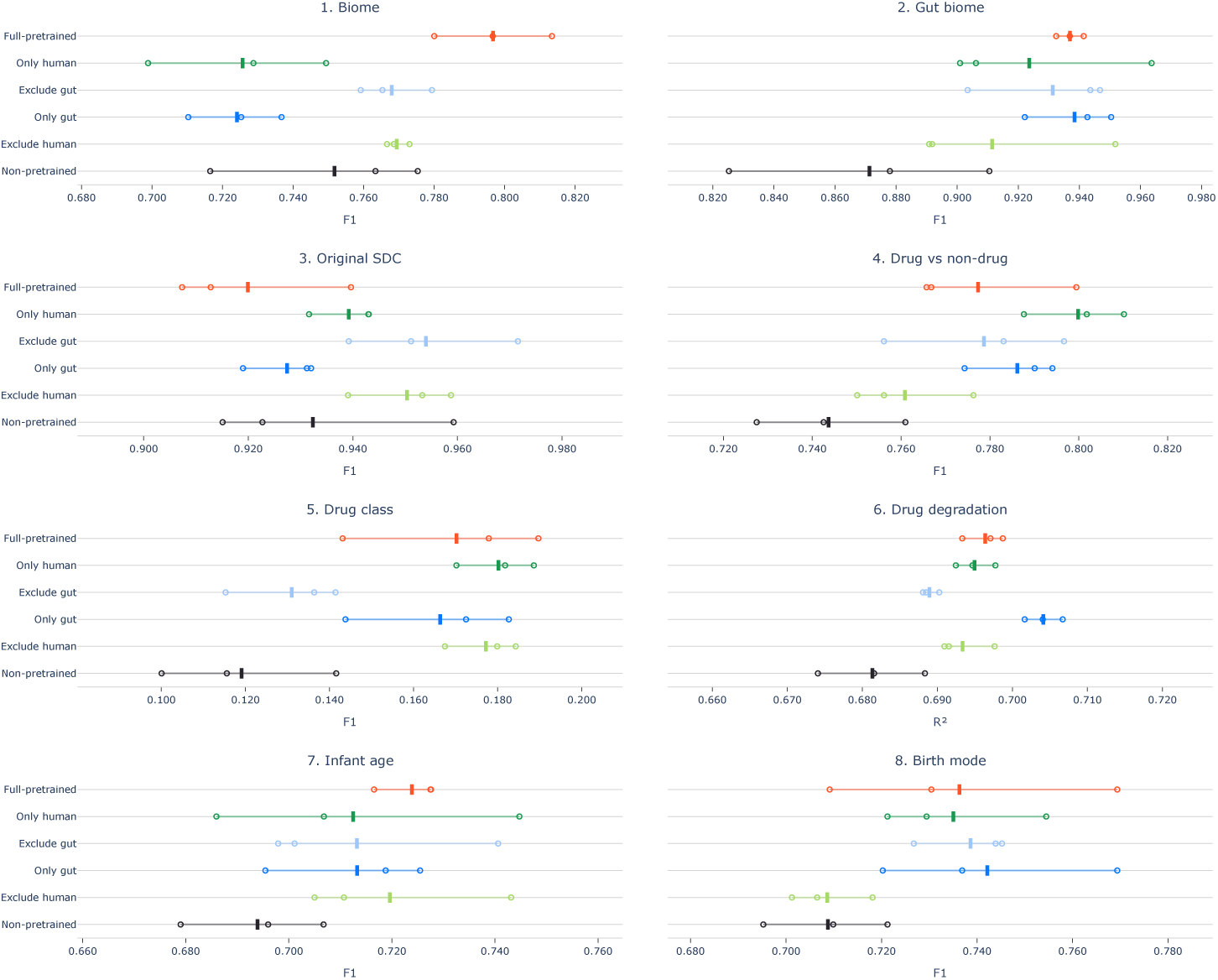
Per-task Compass scores for the 6M model pretrained on different biome subsets of Atlas where all datasets are subsampled down to 110k data points. Classification tasks (1–5, 7–8) are scored by macro-averaged *F*_1_ and the regression task (6) by *R*^2^. Solid lines indicate the mean and hollow circles individual replicates. The cross-biome transfer observed in the aggregate holds across individual tasks.

